# Single-sample proteome enrichment enables missing protein recovery and phenotype association

**DOI:** 10.1101/2021.11.13.468488

**Authors:** Bertrand Jern Han Wong, Weijia Kong, Wilson Wen Bin Goh

## Abstract

Proteomic studies characterize the protein composition of complex biological samples. Despite recent developments in mass spectrometry instrumentation and computational tools, low proteome coverage remains a challenge. To address this, we present Proteome Support Vector Enrichment (PROSE), a fast, scalable, and effective pipeline for scoring protein identifications based on gene co-expression matrices. Using a simple set of observed proteins as input, PROSE gauges the relative importance of proteins in the phenotype. The resultant enrichment scores are interpretable and stable, corresponding well to the source phenotype, thus enabling reproducible recovery of missing proteins. We further demonstrate its utility via reanalysis of the Cancer Cell Line Encyclopedia (CCLE) proteomic data, with prediction of oncogenic dependencies and identification of well-defined regulatory modules. PROSE is available as a user-friendly Python module from https://github.com/bwbio/PROSE.

## Introduction

Proteomics, the high-throughput analysis of proteins in a sample, enables deep study of functional factors underlying a biological condition or phenotype^1^. Liquid chromatography-tandem mass spectrometry (LC-MS/MS) is the most widely used technology in proteomics, supporting the rapid and parallelizable analysis of complex biological samples^1^. In a typical proteomic experiment^1,2^, sample proteins are first digested using a site-specific protease such as trypsin, followed by column-based separation of the resultant peptides, and spectral profiling at the MS1 and/or MS2 levels. Acquired spectra are then compared against theoretical spectra to obtain peptide-spectrum matches (PSMs). Confident PSMs, corresponding to component peptides of full-length proteins, are then used to determine and quantify present proteins in each sample. Proteomics is a rapidly developing technology: The past decade has seen a shift from traditional shotgun proteomic technologies, based on the Data-Dependent Acquisition (DDA) paradigm towards Data-Independent Acquisition (DIA)^3^. In DDA, few precursor ion peaks in MS1 are captured and analyzed in MS2^4^, whereas in DIA, extensively multiplexed MS1 peaks are used to generate complex MS2 signatures, which are then computationally deconvoluted to identify peptides^3,5–7^. This approach allows exhaustive capture of peptide signal but is computationally more complex. Newer instrumental approaches, such as diaPASEF/timsTOF^8^, further expand on ion discrimination and spectral capture capabilities.

Despite these advances, missing proteins are still abundant in the majority of published proteomic datasets^9^. This issue hinders proper downstream analysis, and accurate imputation of these missing values, especially for proteins missing in all replicates, has proven challenging^9,10^. Some authors suggest that this is the result of a computational bottleneck: certain peptides are inherently difficult to identify from their spectra, possibly due to extensive post-translational modifications (PTMs)^11,12^; the identification of modified peptides from the whole proteome entails an untenable increase in search space, with massive computational expense and high risk of false positive matches^12^. The recent development of some tools, such as DIA-NN^13^, have demonstrated some success in identifying these modified peptides. Nevertheless, there remains no clear consensus on how to approach these issues of peptide detectability and proteome coverage beyond improving spectral capture. Furthermore, developing and deploying more sophisticated LC-MS^(n)^ instrumentation may not always be feasible due to high capital/operational costs.

We also need to consider what to do with all the data acquired thus far. Analytics may hold the key towards improving data reuse via data integration and better contextual reasoning. One promising alternative direction is the use of biological context and networks to infer the presence of missing proteins^14,15^. Our recent algorithm PROTREC^15^, which uses protein complexes as contextual information, was able to successfully recover missing proteins across replicates of HeLa lysates and solid cancer samples, using only protein lists as input. In principle, methods such as PROTREC present certain key advantages: they can be adapted to any instrumentation, are independent of spectra, and can be applied to re-analysis of public proteomic datasets, including low-coverage DDA. Given such potential, developing more powerful and wider-scoping techniques that leverage biological context may prove invaluable. PROTREC also employs direct probabilistic reasoning, thus intuitively capturing direct information with high model interpretability. However, machine learning/deep learning approaches with more complex architecture are rapidly gaining popularity, stemming from their ability to capture fuzzy, non-linear or subtle relationships while simultaneously integrating diverse sources of data.

Co-expression networks and matrices, which encode relationships between individual genes, are an interesting and valuable source of biological context; a large body of literature^16^ demonstrates their diverse utility across various applications, including prediction of disease-linked genes^17,18^, functional annotation^19^, and disease state classification^20^. However, to our knowledge, machine learning approaches have yet to be applied to co-expression data for either missing protein recovery or phenotypic enrichment scoring of individual proteins/genes.

In this vein, we have developed Proteome Support Vector Enrichment (PROSE), a machine learning approach based on support vector machines, for missing protein recovery and phenotype association. A simplified schematic of the workflow is presented in **Figure 1a**. PROSE takes a simple list of identified (and unidentified) proteins in a sample, before applying a supervised learning approach on a separate gene co-expression matrix, derived from public RNA-Seq data. PROSE outputs an enrichment score for each protein, based on their association to the putative sample phenotype. These enrichment scores can then be used to infer missing proteins in the dataset, or used for functional analysis. The input protein lists can be derived from a variety of sources, including peptide identifications or relative quantitation data, allowing for flexible analyses. PROSE is made available as a user-friendly Python module.

**Figure 1.**
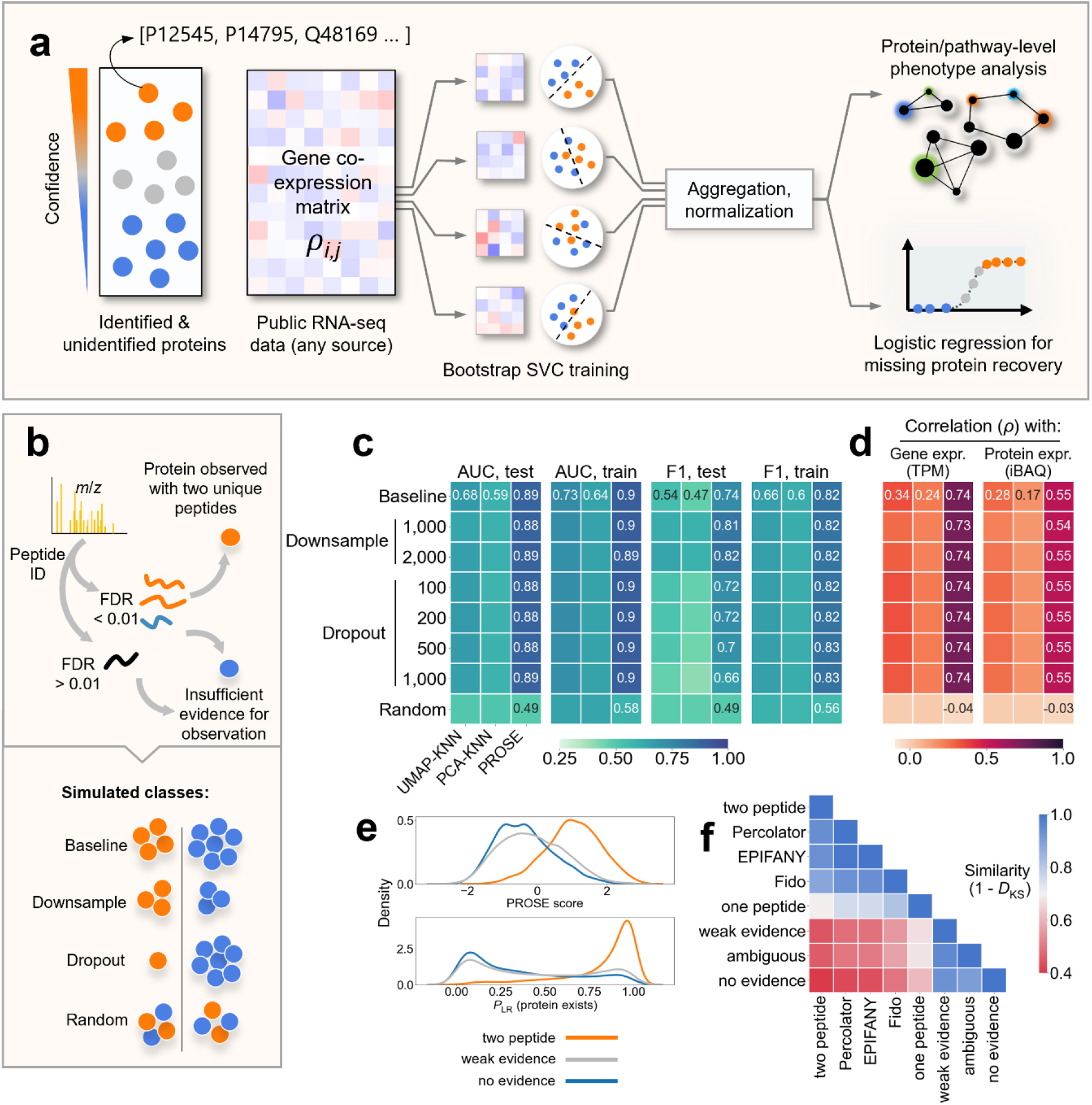
PROSE accurately discriminates between existing and non-existing proteins within a sample. **a**: Schematic representation of PROSE scoring workflow. **b**: Schematic representation of proteome assembly for the HeLa DDA dataset, with simulated missing values for benchmarking. **c**: Comparative performance of PROSE against dimensionality reduction methods for missing protein recovery. **d**: Spearman correlation coefficients *ρ* between each method’s decision function and HeLa RNA-seq data (gene expression; TPM), as well as quantitative proteomics (protein expression; iBAQ). Metrics shown are averaged across 10 simulations, across each of the 4 HeLa DDA replicates. **e**: Representative density estimates of PROSE score and logistic regression probability *P*_LR_ (protein exists) from HeLa R1, demonstrating strong class discrimination. **f**: Similarity of PROSE score distributions between proteins identified under different inference rules, represented by the complement of the Kolmogorov-Smirnov distance (1 − *D_KS_*).

## Results

### PROSE scores are biologically relevant and enable missing protein recovery

We evaluated the performance of PROSE by labelling identified and unidentified proteins derived from a PSM search on HeLa digest spectra **(Fig. 1b)**. We benchmarked PROSE against two baseline *k*-nearest neighbour (KNN) classifiers applied to either PCA or UMAP transformations of the same co-expression matrix **(Fig. 1c)**. In both the baseline and simulated datasets, PROSE showed markedly higher performance in terms of area under the receiver operating characteristic (AUC) and F1 scores, over the baseline classifiers. PROSE scores are also stable between and within HeLa replicates at various bootstrap estimator sizes **(Supplementary Fig. 1)**. The low performance of PCA-KNN and UMAP-KNN is likely due to the non-linear separability of the PCA/UMAP transformations **(Supplementary Fig. 2)**, highlighting the necessity of the machine learning approach. Interestingly, UMAP-KNN performs better than random, stemming from the fact that several proteins with similar expression profiles cluster together on the UMAP, leading to a preference for low *k* during KNN optimization.

To assess the biological relevance of each scoring approach, we considered the Spearman correlation between the score vector and the related RNA-Seq or quantitative proteomic data for HeLa **(Fig. 1d)**. PROSE scores show strong positive correlation with transcript per million (TPM; *ρ* = 0.74) and modest correlation with protein intensity-based absolute quantitation (iBAQ; *ρ* = 0.55), while both KNN models provide weak correlations.

The distinct PROSE score distributions enable high- and low-confidence protein identifications **(Fig. 1e)**, where a positive class probability *P*_LR_ > 0.5 is considered as an identified protein, enable strong separation of positive and negative classes. Further comparisons of score distributions of protein sets generated under different inference rules **(Fig. 1f)** highlight the similar distribution of proteins supported only ambiguous peptides and those with no support, suggesting that a minority of these proteins occur in the sample.

To examine if PROSE scores show tissue-specific effects, we obtained correlations between the HeLa scores and TPM values of various cell line **(Supplementary Fig. 3a)**. In addition to strong correlation to HeLa, we also observed strong correlation to cell lines of similar origin, in particular those from epithelial lineages. Interestingly, the originating tissues with the largest correlation generally stem from those of similar mucosal nature, including the chrodate pharynx, oral cavity, and urinary bladder. The correlation was lowest to blood/lymphoid tissues, which highlight their distinct transcriptomic niche^10,21^. Scatterplot comparisons of PROSE scores to either HeLa or EJM (plasma cell/multiple myeloma line) show that proteins with high PROSE scores have high TPM in HeLa but not necessarily in EJM **(Supplementary Fig. 3b)**, whereas proteins lacking TPM-level evidence generally have low PROSE scores, but few are recovered in HeLa.

### Scores of Cancer Cell Line Encyclopedia (CCLE) samples correlate with tissue of origin and gene dependency

The CCLE proteome data comprises protein expression levels, obtained using a tandem mass tag (TMT; TMT quants) approach^10^. This poses a challenge in interpretation, as these values are strictly relative and not normalized across the entire dataset^22^. Phenotypic scoring approaches such as PROSE may provide a means of transforming these quants into meaningful intra-cell line distributions. We applied PROSE to a set of positive and negative proteins in each cell line, defined by a heuristic split of quantified proteins by the median expression value (relative quants across a multiplex; TMT quant). For the vast majority of proteins, PROSE scores correlate well with corresponding RNA-Seq data (as TPM) and TMT quants, across the CCLE samples **(Fig. 2a)**. The correlation for scores within a cell line were modest **(Fig. 2b)** but largely positive, although there was a noticeable increase in the mean correlation between PROSE score and TMT quant compared to between TMT and TPM. Subsequent cluster analysis via UMAP **(Fig. 2c**) and correlation analysis of matched score distributions, grouped by tissue of origin **(Fig. 2d-e)**, demonstrate successful recovery of tissue-specific effects. In addition, we predicted the cell-line specific relevance of certain proteins with transcriptomic evidence that have not been previously observed **(Supplementary Fig. 4a,b)** in vivo, in either proteomic or single-protein studies, though in general they carry low mean PROSE scores across cell lines.

**Figure 2.**
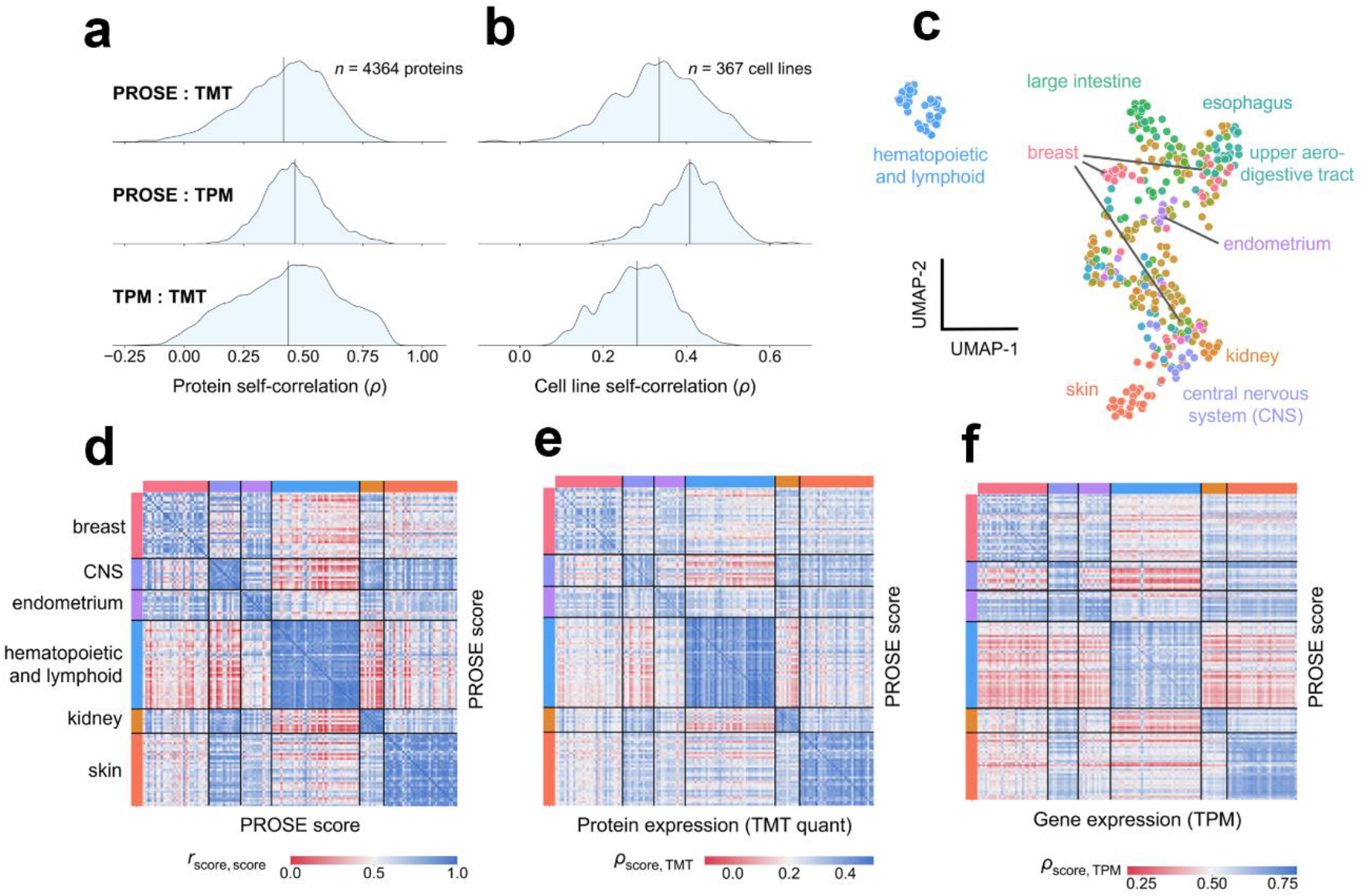
PROSE scores show specificity for tissue of origin. **a**: Distribution of Spearman correlations coefficients *ρ* for each protein, obtained by comparing the PROSE, TMT (protein expression), or TPM (gene expression) vectors. **b**: Distribution of *ρ* for each cell line, obtained similarly. Vertical lines represent mean values. **c**: UMAP derived from PROSE scores highlighting tissue clustering. **d-f**: Heatmaps demonstrating tissue-specific correlation between PROSE scores, TMT, and TPM. A select subset of tissues are shown; blood (hematopoietic and lymphoid tissues) is noticeably dissimilar to most non-blood cell lines. *r*, Pearson correlation coefficient; *ρ*, Spearman correlation coefficient.

To further evaluate the biological relevance of these scores within a sample, we compared the score distributions of high- and low-dependency genes within cell lines, using the dependency data generated from the DRIVE high-throughput deep RNAi screen^23,24^. For simplicity, high-dependency genes were defined as those in the 95^th^ percentile by dependency (essentially, the negative value of the DEMETER2-corrected shRNA abundance^24^), with the rest considered low-dependency. Compared to TMT quants, PROSE scores enabled effective separation of these groups in the majority of cell lines, using either the Kolmogorov-Smirnoff statistic **(Fig. 3a,b)** or difference of medians **(Fig. 3c,d)** as the distance metric. The concentration of high-dependency genes in the high score region was visualized for select cell lines **(Fig. 3e)**. As TMT quants correspond to relative expression of a protein within the multiplex, there is a reasonable expectation that high quants may correlate to increased dependency (i.e. in the case of oncogene addiction). However, no intra-cell line visual separation is observed when using the normalized TMT quants **(Fig. 3f)**, although inter-cell line dependency for individual genes is known to be correlated to TMT quants. Interestingly, the lack of separability by TMT was observed despite a positive correlation between the score and TMT quant within cell lines **(Supplementary Fig. 4c)**. Overall, our findings demonstrate the utility of PROSE to generate meaningful intra-sample information from inter-sample quantitation.

**Figure 3.**
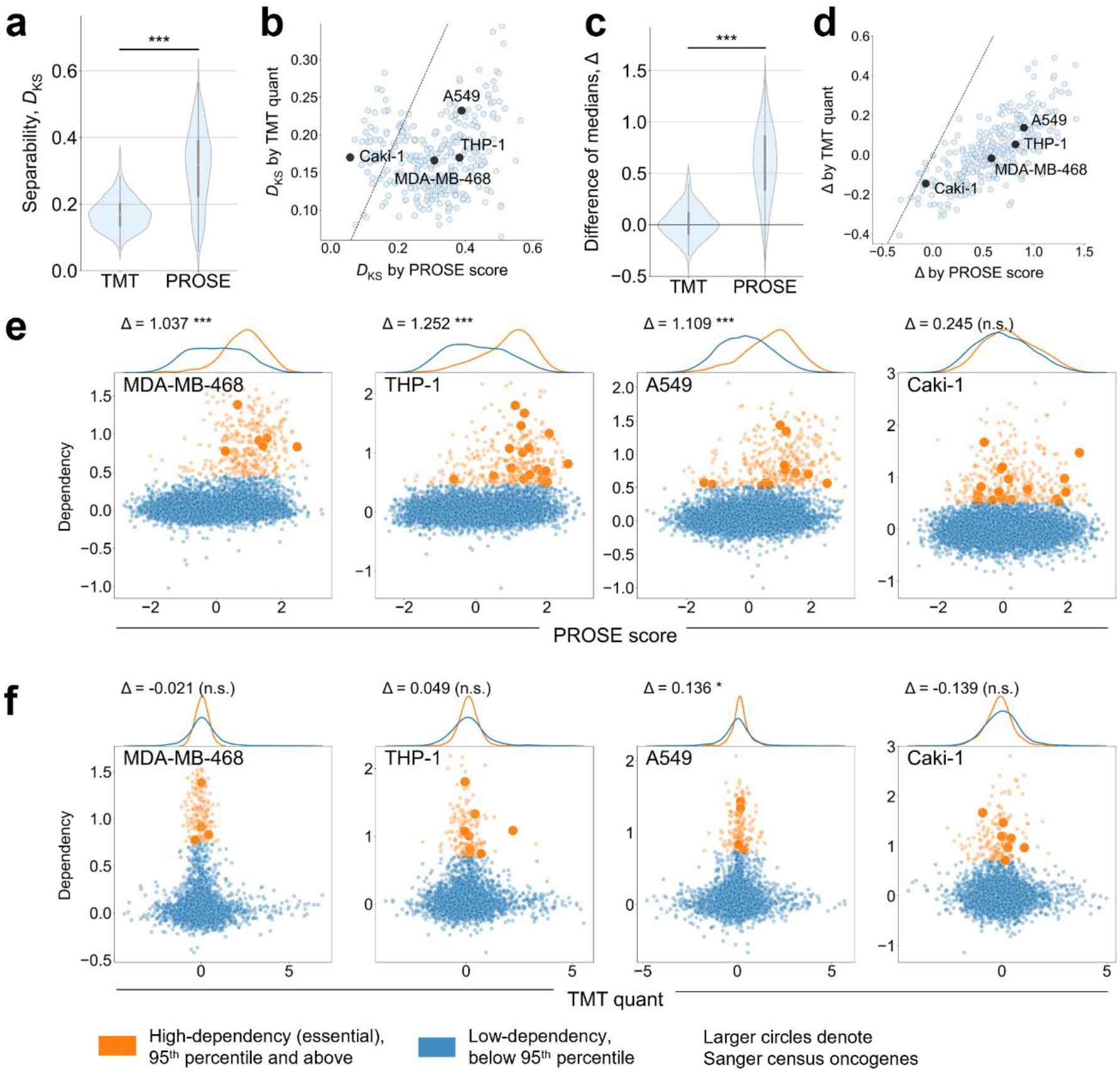
Essential genes have high PROSE scores. For each CCLE cell line, the set of genes were split into high-dependency (95^th^ percentile and above) and low-dependency (below 95^th^ percentile) subsets. **a, b**: Separability of subsets measured by the Kolmogorov-Smirnov distance metric (*D*_KS_). **c, d**: Separability measured by the difference of medians (high-dependency – low-dependency; Δ). Mann-Whitney U test (\*\*\**p* < 0.001). For scatterplots, each point represents an individual cell line. The dashed line indicates no difference between separation by PROSE or TMT. For comparisons, each distribution was Z-normalized. **e**: Representative cell lines highlighting the skewed PROSE score distribution of high-dependency genes. **f**: Representative cell lines highlighting the lack of separability using TMT quant. Each point represents an individual gene/corresponding protein. Several proteins, both high- and low-dependency, were missing from the proteomic screen and hence excluded from analysis. Bootstrap test for Δ > 0, with 10^4^ randomizations (\**p* < 0.05, \*\*\**p* < 0.001, n.s = not significant).

### PROSE-FastICA identifies compact co-regulated modules across CCLE samples

To determine if PROSE scores could be used to generate meaningful co-regulation modules (i.e. protein sets with well-defined expression patterns) across the CCLE cell lines, we applied a FastICA-FDR-like algorithm^25,26^ to decompose the score matrix into independent signal vectors containing significant proteins **(Fig. 4a)**. Of the 200 components tested, using a significance threshold of 10^−6^ for protein filtering, we were able to extract 114 significant, minimally overlapping modules of sizes between 10 and 177.

**Figure 4.**
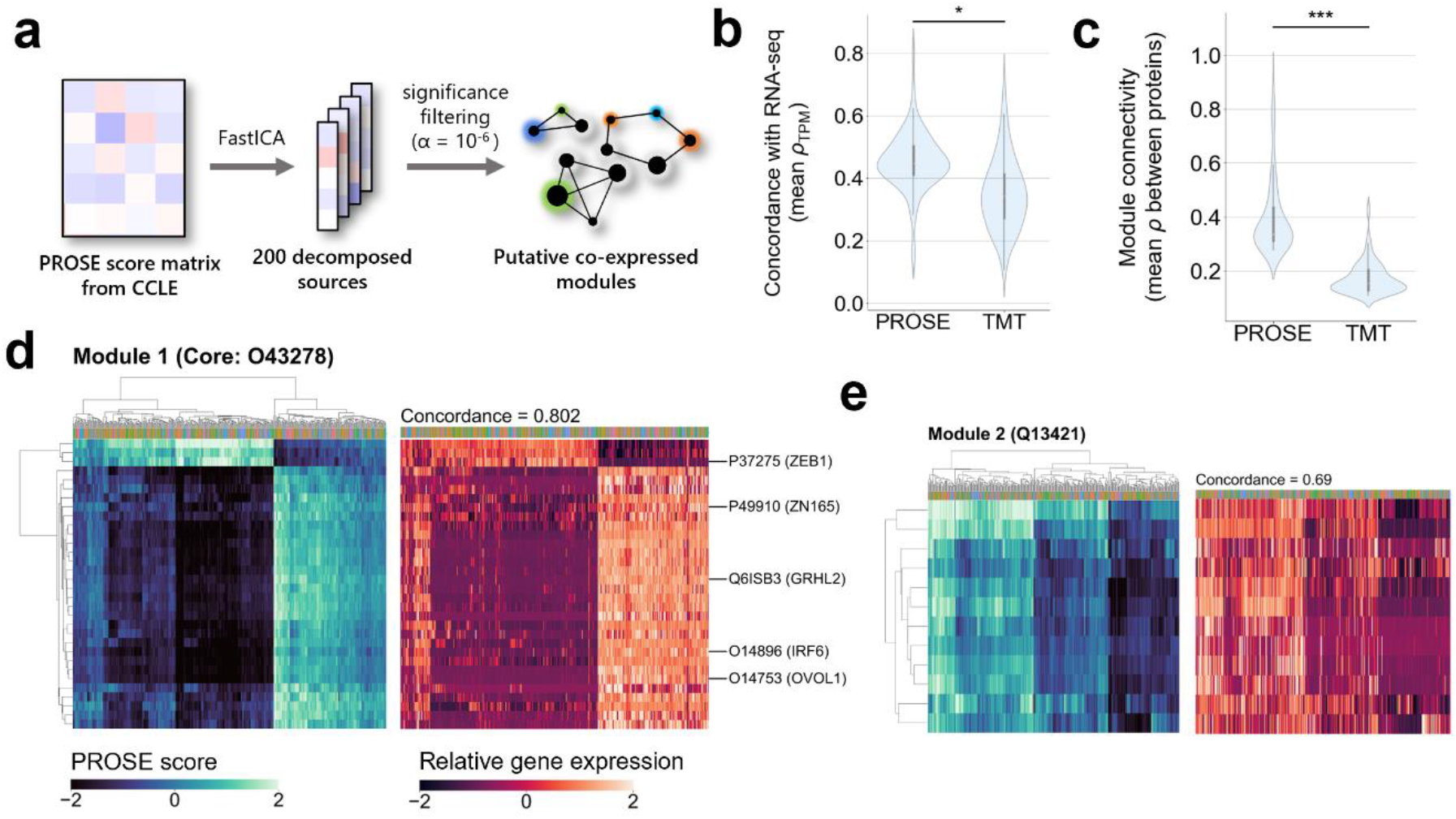
PROSE scores enable coherent module discovery from protein lists. **a**: Module discovery approach by FastICA. **b, c**: Comparison of performance metrics for modules determined from either PROSE or TMT with FastICA. Modules discovered with PROSE correlate more strongly to RNA-seq data, and show better internal connectivity, compared to those discovered from TMT quants. Bootstrap test with 10^5^ randomizations (\**p* < 0.05, \*\*\**p* < 0.001). **d**: Highest-concordance module discovered by PROSE-FastICA, identifying a *ZEB1-IRF6-OVOL1* co-regulatory system, implicated in epithelial-mesenchymal transition. Known transcription factors are annotated on the left. **e**: Second highest-concordance module showing similar clustering.

Modules defined from the score matrix showed higher concordance with the underlying RNA-Seq data **(Fig. 4b)**, as well as intra-module connectivity **(Fig. 4c)**, as compared to modules from the TMT quant matrix. The highest-concordance module was defined by the transcription factors ZEB1^27–29^, ZN165^30,31^, GRHL2^32^, IRF6^28^, and OVOL1^33^, which have known roles in the regulation of epithelial-mesenchymal transition (EMT) and related pathways, which underlies metastatic potential. We observed a striking similarity between the scores and corresponding gene expression within the module **(Fig. 4d)**, with no observed tissue-specific effects. Similar visual similarities were observed for other modules **(Fig. 4e; Supplementary Fig. 5)**. In addition, we were able to recover well-defined modules with complex clustering patterns. Of note, one module was comprised almost entirely of the HOXA-HOXB homeobox transcription factors, which are involved in regulation of developmental processes^34^ **(Supplementary Fig. 6)**, which did not cluster strongly by cell line.

## Discussion

In this study, we have developed PROSE as a technique that leverages on the biological and phenotypic information contained in public transcriptomic datasets. Transforming this quantitative data into co-expression matrices allowed us to score individual proteins by their propensity for upregulation in an expression-naïve manner.

At present, much co-expression research focuses on understanding and harnessing the properties of information transfer between elements in a network (for instance, in weighted gene co-expression network analysis^35,36^); sophisticated machine learning models on *de novo* co-expression networks have also seen success in the imputation of missing values, including graph-based methods for predicting expression in single-cell RNA-Seq datasets^37^. Nevertheless, our findings suggest that simpler, topology-independent models may be sufficient to capture diverse expression patterns. In our case, the use of a co-expression matrix as a feature space is perhaps analogous to the two-class regression model on first neighbours in a co-expression network. Despite this, we find it difficult to achieve a satisfying separation of the observed and unobserved genes via dimensionality reduction approaches (see **Supplementary Fig. 2**), which suggests that expression can only be predicted by higher-order relationships between co-regulated genes. Considering the high performance that can be achieved from the co-expression matrix alone, further investigation into its useful properties and pitfalls is warranted: indeed, we are aware that similarities and correlations have powerful properties in many key bioinformatics application areas (sequence comparisons and function predictions being notable examples), but can also lead towards undesirable outcomes such as doppelgänger effects^38^.

Through rescoring of the CCLE proteomic data, we show that relative protein expression across cell lines can be transformed into a meaningful intra-sample distribution that reflects biological features, including gene dependency. Importantly, we were able to generate scores for missing proteins in each multiplex, many of which are encoded by high-dependency genes. In this manner, the resultant scores may be complementary to imputation methods, which are generally complex to apply to relative quantitation data^39^ and in some cases, depend on the presence of multiple replicates to generate accurate expression distributions^9^. As PROSE scores show clear relationships to the underlying gene and protein expression, they may possibly be used as additional feature to improve imputation accuracy.

In conclusion, our results encourage further study into methods that can effectively capitalize on the information value of co-expression matrices. We find that models such as PROSE for protein enrichment may provide a useful means of transforming complex proteomic data into interpretable scores. While we have focused on PROSE as a tool for missing protein recovery using protein lists, in principle it can also be used to generate enrichment scores from gene/protein sets generated from any number of absolute or relative quantitation paradigms, providing a high degree of usage flexibility.

## Materials and Methods

### Gene co-expression matrix construction

The Klijn et al. (2015) dataset^40^, comprising RNA-Seq results for 675 common cancer cell lines, was downloaded in TPM format from ArrayExpress^41^ (https://www.ebi.ac.uk/arrayexpress/) under the accession number E-MTAB-2706. Gene identifiers were converted to UniProt IDs, with only protein-coding genes retained. Genes with consistently low expression (median TPM < 1) were excluded. The co-expression matrix was obtained by calculating the Spearman correlation coefficient *ρ* between each pair of genes. To limit the feature vector to a tight set of panel genes, we selected those with high variability in expression and co-regulation. These were defined as the top 25% of genes ranked by coefficient of variation (CV; of TPM), as well as standard deviation of correlation coefficients, yielding 1216 panel genes **(Supplementary Fig. 7a)**. We verified that the panel genes enabled differentiation and clustering of cell lines by tissue origin by visual inspection of UMAP plots and heatmaps of gene expression profiles **(Supplementary Fig. 7b-d)**. The resultant 17054 × 1216 matrix is an array of protein identifiers with associated Spearman correlations to each panel gene. For our analyses, TPM values were log2 transformed using a pseudocount of 1.

### PROSE

The core of the PROSE module is built from the *scikit.learn* Python library^42^. Using a list of positive (observed) proteins, negative (unobserved) proteins, and the gene co-expression matrix above, PROSE returns a table containing a list of these proteins, their associated PROSE scores, predicted labels, and their logistic regression probabilities.

Briefly, for a given proteomic screen, we define these observed proteins as a set of high-confidence protein identifications. Typically, the two-peptide rule with a peptide FDR threshold of 0.01 was applied. The set of unobserved proteins is defined as those with low (FDR > 0.3) or no peptide evidence. In general, the sizes of these class are unlikely to be balanced, hence synthetic minority oversampling (SMOTE)^43^ is applied to balance the observations. An ensemble of linear support vector machines is applied to generate hyperplanes separating the positive and null observations based on their correlation to the panel genes. To enhance stability and minimize computational expense, a bootstrap-aggregating (bagging)^44^ approach was used to generate a series of weak learners that converge upon correct estimates for phenotypic relevance. In our implementation, *k* sub-classifiers are trained on a bootstrap sample of 100 proteins drawn from each class, under 50 panel genes. Unless otherwise stated, in our analyses *k* = 1000 bootstrap samples were used. The use of bagging renders PROSE scalable to any number of panel genes and observations, for instance in an expanded transcript-level co-expression network. However, it may be necessary to increase *k* to improve coverage of relevant features as necessary. Alternatively, a similar panel gene selection approach can be applied to maximize coverage of the highest-variance features.

Formally, the decision function *g*(*i*) for a given protein *i* is defined as the mean signed distance between the hyperplanes and the point of interest:

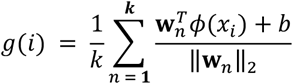

Where *g*(*i*) > 0 represents the positive class and g(*i*) < 0 represents the null class. We avoid directly making class predictions on *g*(*i*) due to bias towards the null class amplified by the bagging strategy. To further mitigate imbalanced class effects, we instead consider the Z-normalized metric, yielding the final PROSE enrichment score:

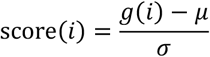

Where *μ* and *σ* are the mean and standard deviation over the set of all tested proteins in the co-expression matrix. The ranked output score vector is generally sufficient for functional analyses of phenotype. However, for the purpose of missing protein recovery, we apply logistic regression on score(*i*) for class prediction, where protein probability *P*_LR_ was estimated via 5-fold cross validation. Classifier performance was evaluated on a stratified set of 200 holdout proteins. For correlation analyses between the HeLa PROSE scores and gene expression, the TPM data from Klijn et al. (2015) above was used. For correlation analyses to HeLa protein expression, the HeLa proteomic dataset from Bekker-Jensen et al. (2017) was used^45^. The processed protein expression data as iBAQ was downloaded from https://doi.org/10.1016/j.cels.2017.05.009.

### *k*-nearest neighbour (KNN) classification

The co-expression matrix used for PROSE analysis was transformed either by PCA or UMAP, retaining the top 2 components as KNN inputs (see **Supplementary Fig. 2**). For the PCA-transformed matrix, each component was scaled to its explained variance. The set of positive and negative protein observations were used as class labels. For each instance, a grid search was performed to optimize *k*, which was allowed to vary between 5 and 50. Classifier performance was evaluated on a set of 200 holdout proteins stratified by the provided classes.

### Raw proteomic data processing

The Mehta et al. (2021) HeLa DDA dataset^46^ comprises commercial HeLa digest standards in quadruplicate, acquired on a Orbitrap Fusion Lumos Tribrid MS instrument. Data was downloaded from the PRIDE database^37^ (https://www.ebi.ac.uk/pride/) with the accession number PXD022448, and processed using OpenMS^47,48^. For each replicate, raw files were converted to .mzml files. PSMs for tryptic peptides of length 6 to 40 inclusive were identified using the MS-GF+ search engine with a precursor mass tolerance of 10.0 ppm. 1 missed cleavage was allowed. Carbamidomethyl (C) was set as the fixed modification and oxidation (M) was set as a variable modification. Peptide-level FDR was estimated with a conventional target-decoy approach; the target-decoy FASTA database supplied to MS-GF+ was constructed from the set of verified UniProt sequences by concatenation of the reversed sequences. Proteins were subsequently assembled directly from the set of peptide identifications, or alternatively inferred using the Percolator^49^, Fido^50^ or EPIFANY^51^ engines. For EPIFANY inference, peptides were first rescored by Percolator. Default parameters were used for all search and identity engines except where highlighted. The inferential pipelines used for identifying positive and null protein identifications for this proteome assembly are summarized in **Supplementary Table 1**.

### Cancer Cell Line Encylopedia (CCLE) analyses

PROSE scoring was performed on the CCLE proteomic dataset from Nusinow et al. (2020)^10^. This dataset comprises normalized tandem-mass-tag (TMT) quantitation of proteomes across multiplexed CCLE samples acquired on either a Orbitrap Fusion or Orbitrap Fusion Lumos MS instrument. For a given cell line, protein identifications were obtained via parsimonious assembly within the multiplexed sample without imputation, and the assembly displays a wide number of missing proteins. Normalized protein quants were downloaded from https://gygi.hms.harvard.edu/publications/ccle.html. Only SwissProt (reviewed) proteins were considered in our analysis. Due to the computational expense of reprocessing a large volume of raw proteomic data, we instead opted to generate heuristic sets from the processed data; the positive and null classes were obtained by equally partitioning the set of proteins based on their intra-sample TMT quants. Note that while these quants were previously found to correlate to gene expression levels, they specifically measure protein expression relative to other samples in the multiplex.

For correlation analysis to RNA-Seq, we used the mRNA expression data from Barretina et al. (2012)^21^. Data was downloaded from ArrayExpress^41^ under the accession number E-MTAB-2770. Further correlation analyses were performed on the DRIVE shRNA data from McDonald et al. (2017)^23^, which comprises deep RNAi screens for gene dependencies in 398 cancer cell lines. We used DEMETER2-adjusted scores from McFarland et al (2018) as highly accurate estimates of gene dependency^24^. Data was downloaded from https://figshare.com/articles/dataset/DEMETER2_data/6025238. For simplicity, gene dependence is defined as the negative of the DEMETER2 score (higher dependence indicates essentiality). Oncogenes and proto-oncogenes were annotated using the Sanger COSMIC Cancer Gene Census (CGC)^52^. Our analyses focused on cell lines occurring across these datasets. PROSE scores for the CCLE proteomics dataset are provided as **Supplementary File 1**.

### FastICA for module detection

Co-expressed modules were identified from either the CCLE PROSE score matrix or the TMT quant matrix using an adapted FastICA-FDA approach described previously^25^. The resultant modules for thresholds 10^−4^, 10^−5^, 10^−6^ and raw *p*-values are provided as **Supplementary Files 2-5**.

Briefly, the *scikit.learn* implementation of FastICA was used to decompose each matrix into 200 signals. For each signal vector, we applied a Gaussian fit to generate the null model. Proteins with a probability (of occurrence under the null model) lower than the given threshold were considered module components. For a given module containing *n* proteins, the concordance with the RNA-Seq data was estimated by taking the average of the Spearman correlation for each protein *i*, between the source vector and corresponding TPM vector:

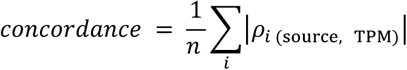

The internal connectivity *C_int_* was estimated by taking the mean of mean Spearman correlations between pairwise proteins *i* and *j* in the source matrix:

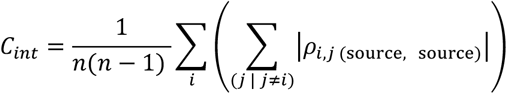

### Gene set overrepresentation analysis (GSOA)

GSOA was performed using the Panther 16.0 webtool^53^ (http://www.pantherdb.org/) to identify GO-Slim (Biological Process) terms associated with PROSE-FastICA modules. Significance was estimated using Fisher’s exact test with FDR correction. Terms with FDR < 0.05 were considered significant.

### Generic analyses

All analyses were performed in a Python 3.8 environment except where otherwise noted. Plotting and visualization was performed using the *matplotlib*^54^ and *seaborn*^55^ libraries. Statistical testing was done with the *scipy*^56^ library. UMAP was performed with the *umap*^57^ library.

## Data Availability

All data not available in the Supplementary Files is available from the authors upon reasonable request. The public datasets used in our analyses are summarized in **Supplementary Table 2**.

## Code Availability

PROSE is available as a Python module from https://github.com/bwbio/PROSE. Code to replicate our analyses is available from https://github.com/bwbio/PROSE/tree/main/analysis_scripts. Note that, as PROSE is built on the *sklearn LinearSVC*, which uses C for fast optimization, the output is not entirely deterministic even when passing seeds to the pseudorandom number generator. We recommend setting the parameter n > 1000 for *bag_kwargs*={‘*n_estimators*’ : *n*}, to produce sufficiently stable inter- and intra-replicate scores, though lower values can be used for aggregate analyses over large gene/protein sets.

## Acknowledgements

This project is supported in part by a Ministry of Education (MOE) AcRF Tier 1 (RG35/20) award to Goh WWB.

## Author contributions

Wong BJH conceptualized PROSE and conducted the analyses. Kong W and Goh WWB provided supervision. Wong BJH, Kong W, and Goh WWB co-wrote the manuscript.

