## Supplementary Information for "Single-sample proteome enrichment enables missing protein recovery and phenotype association"

[^] First author

[*] Corresponding author(s)

**Contents**

**Supplementary Figures**

Supplementary Figure 1. Stability of PROSE scores between HeLa DDA replicates using *k* = 1000 or 100 bootstrapped estimators.

Supplementary Figure 2. PCA projection and UMAP of gene co-expression matrix

Supplementary Figure 3. Visualization of RNA-seq-PROSE score correlations

Supplementary Figure 4. Relationship between TMT quant and PROSE score within cell lines (Related to Figure 4)

Supplementary Figure 5. Additional modules detected by PROSE-FastICA.

Supplementary Figure 6. Module 19 containing the HOXA-HOXB locus

Supplementary Figure 7. Panel gene selection and characterization

**Supplementary Tables**

Supplementary Table 1. Criteria for protein support level classification

Supplementary Table 2. Datasets used in analysis

**Supplementary Files**

Supplementary File 1. PROSE score matrix of CCLE data

Supplementary Files 2-4. PROSE-FastICA modules between sizes 10-1000 defined using a *p*-value threshold of $10^{-4}$, $10^{-5}$, and $10^{-6}$, respectively. Identified modules are sorted by concordance, with the corresponding signal matrix component annotated.

Supplementary File 5. PROSE-FastICA decomposition of the source PROSE score matrix. Values shown are *p*-value transformed FastICA outputs, estimated as the one-tailed probability of containment under the null model.

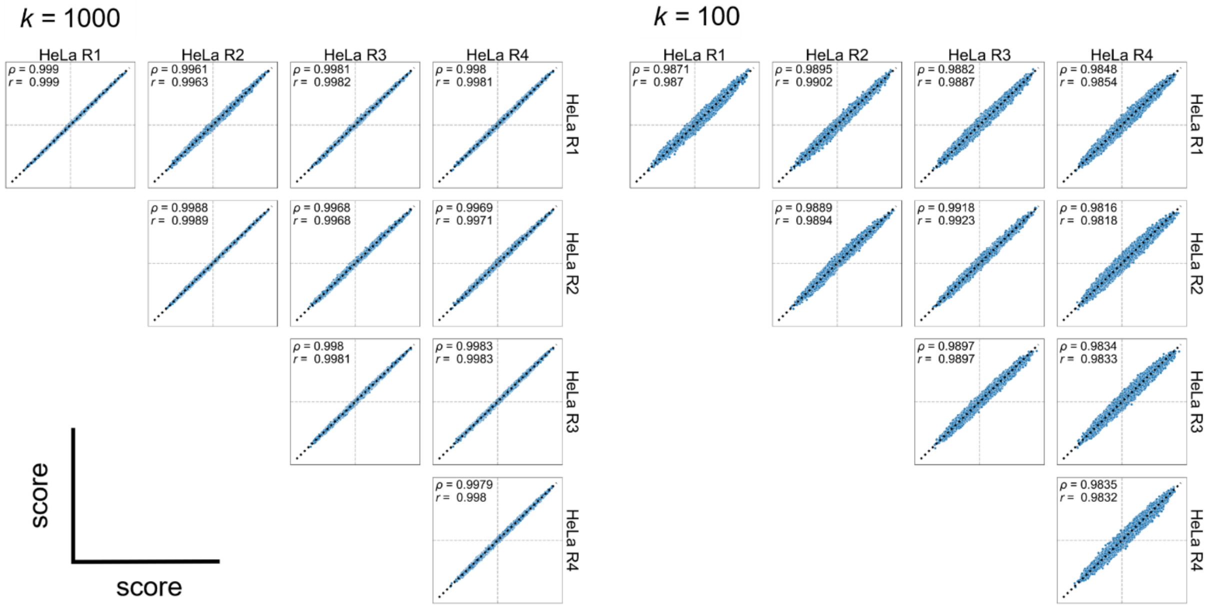


**Supplementary Figure 1. Stability of PROSE scores between HeLa DDA replicates using $k$ = 1000 or 100 bootstrapped estimators.** Each point represents a single protein. Grey dotted lines represent PROSE score = 0 in each axis. Black diagonal dotted line represents score(replicate A) = score(replicate B). Additional values represent Pearson ($r$) and Spearman ($\rho$) correlation coefficients.

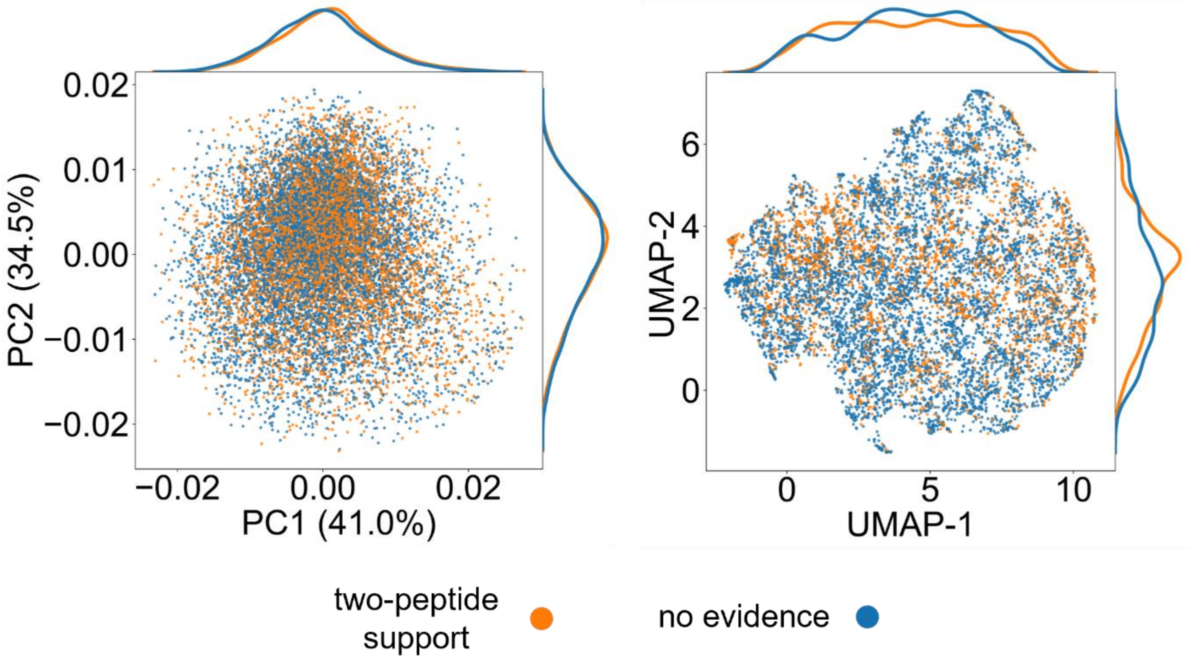


**Supplementary Figure 2. PCA projection and UMAP of gene co-expression matrix.** Each point represents a single protein/associated gene. PCA shows a lack of linear separability, while UMAP shows noticeable but minor clustering of observed/unobserved proteins, leading to better-than-random KNN models.

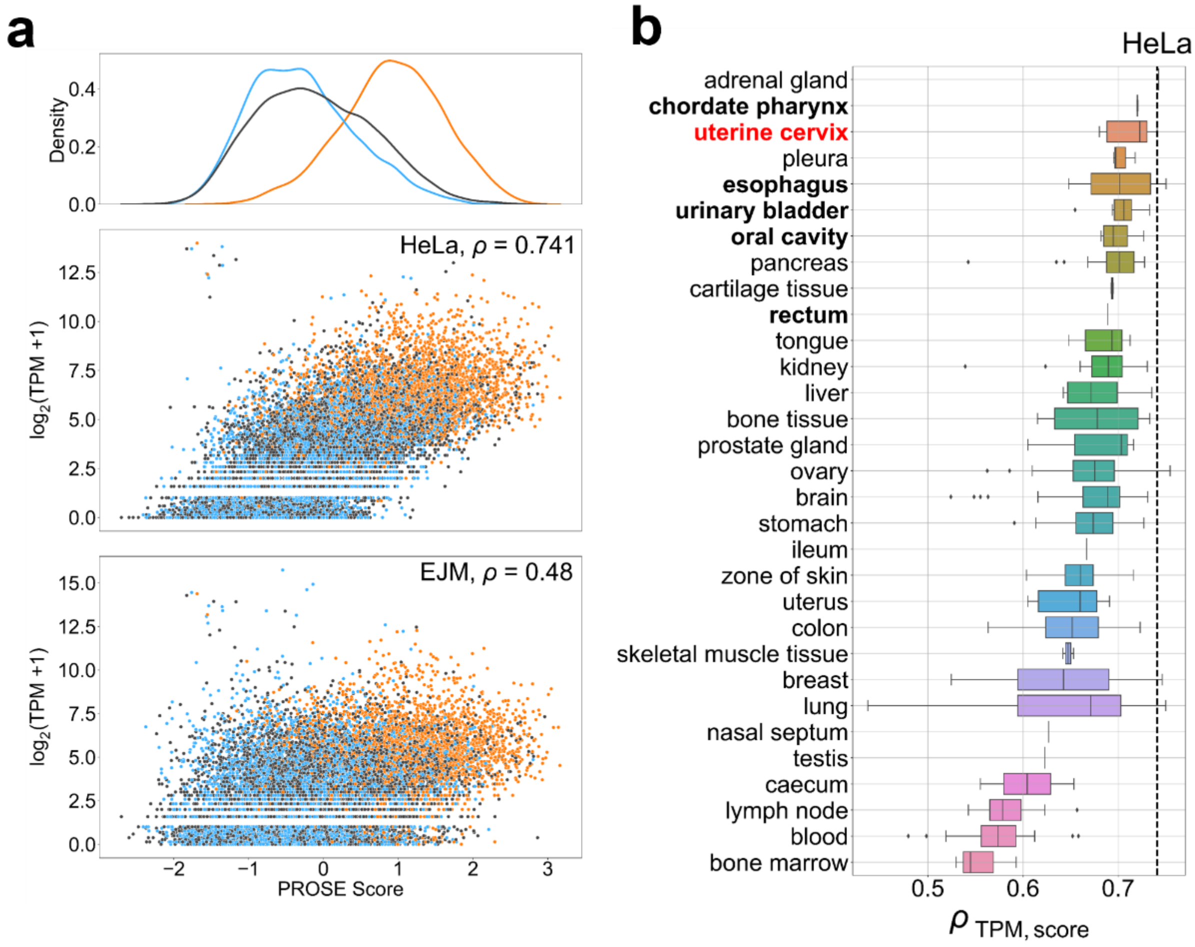


**Supplementary Figure 3. Visualization of RNA-seq-PROSE score correlations. a**: Scatterplot visualization of TPM-PROSE score associations. PROSE scores were obtained from HeLa R1. TPM from HeLa (source) and EJM (unrelated; plasma cell myeloma) cell lines was taken from the Klijn et al. (2015) RNA-seq dataset[1]. b: Correlation of HeLa R1 PROSE scores to TPM in various tissues, sorted by mean value. TPM from HeLa is shown as a dotted line. HeLa is derived from the uterine cervix (bold, in red). Tissues of similar mesodermal origin are bolded. $\rho$, Spearman correlation coefficient.

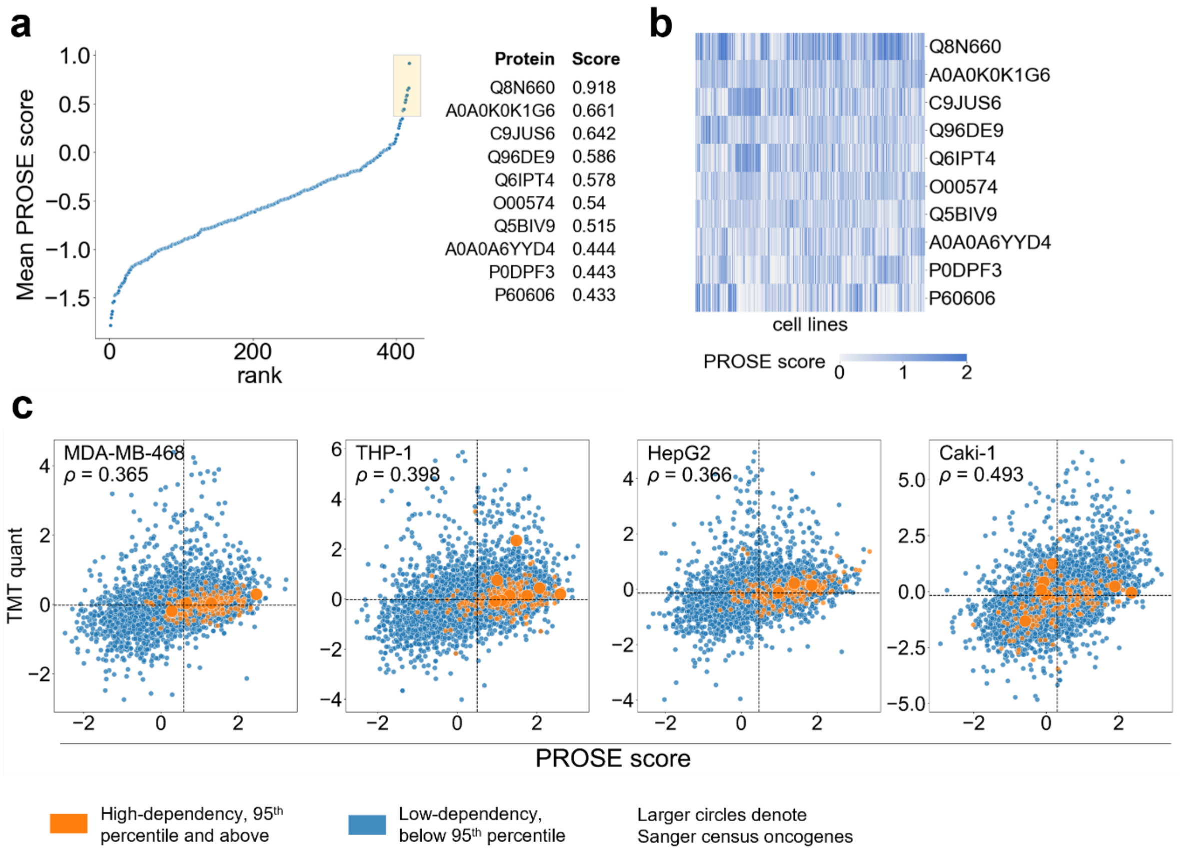


**Supplementary Figure 4. Further analyses into CCLE prose scores. a**: Mean PROSE scores of proteins with only transcript-level evidence according to the Human Proteome Project[1] (HPP; protein evidence level 2 PE2). A majority of PE2 proteins have low PROSE scores. The top 10 proteins by PROSE score are highlighted. **b:** Heatmap of PROSE scores of the top 10 proteins across CCLE samples, suggesting cell-line specific upregulation. Q8N660 is the predicted neuroblastoma breakpoint family protein 15 (NBPF15; alternatively, NBPF16) synthetic fragments reveal domains with amyloid-like characteristic[2], but the endogenous protein is presently undetected. **c:** Scatterplots highlighting the relationship between TMT quant and PROSE score (related to **Fig. 3d,e**). Each point represents a gene/corresponding protein. Vertical and horizontal dotted lines represent the median value for the corresponding variables. For most cell lines, high-dependency genes are concentrated in the high-PROSE score region. In essentially all cases, there is no clear bias for either side of the TMT quant distribution, demonstrating that the relative essentiality of a gene within a cell line does not correspond to its relative protein abundance across cell lines (i.e., as an upregulated protein).

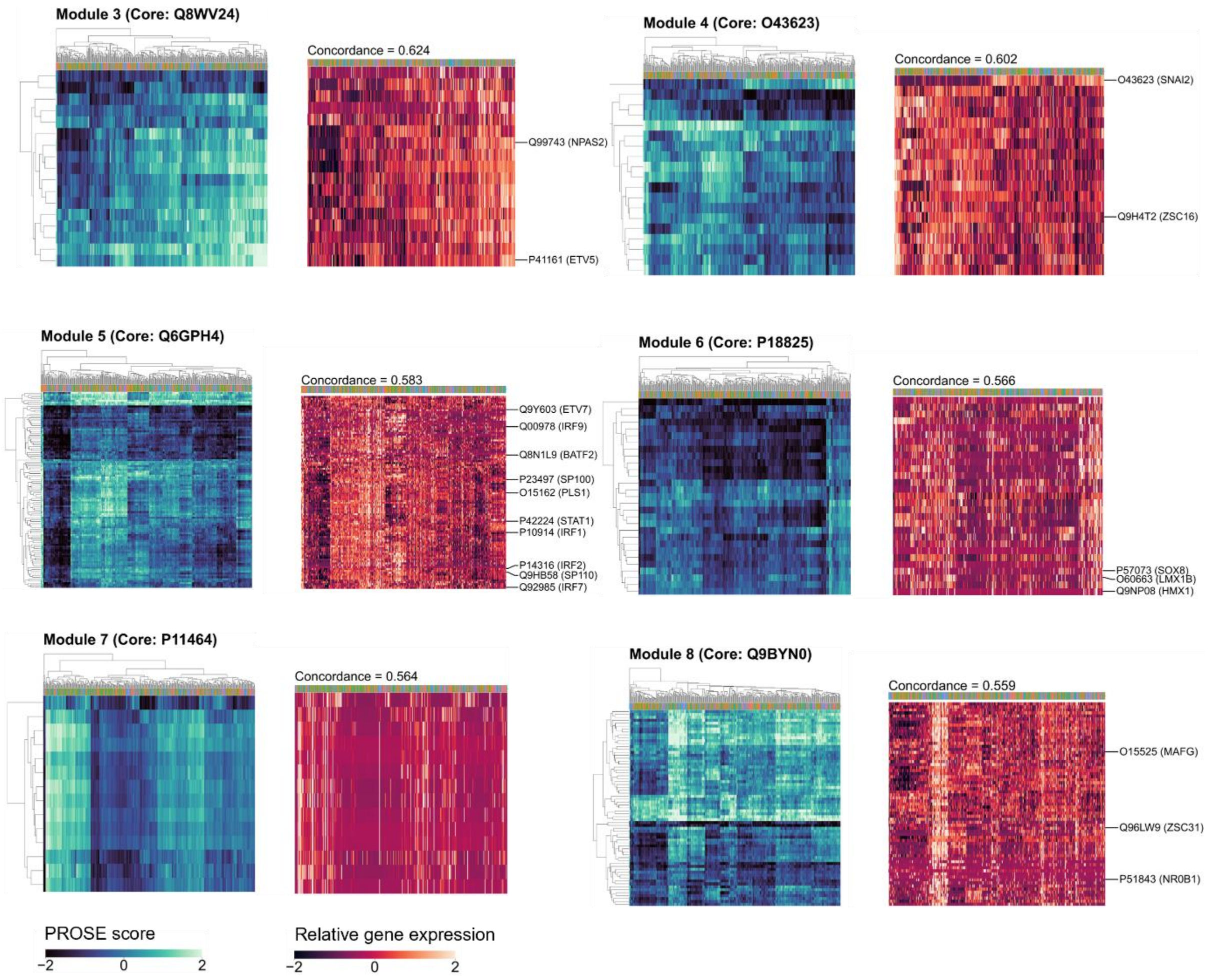


**Supplementary Figure 5. Additional modules detected by PROSE-FastICA:** Modules 3-8 identified using PROSE-FastICA show strong concordance between the clustered score and TPM matrices. Visual inspection reveals higher emphasis on smaller-scale biclusters, rather than global-scale ones as observed for Module 1 (see **Fig. 4d**). Known transcription factors are annotated on the right. The majority of these modules can be described by one or more transcription factors (see **Supplementary Table 1**).

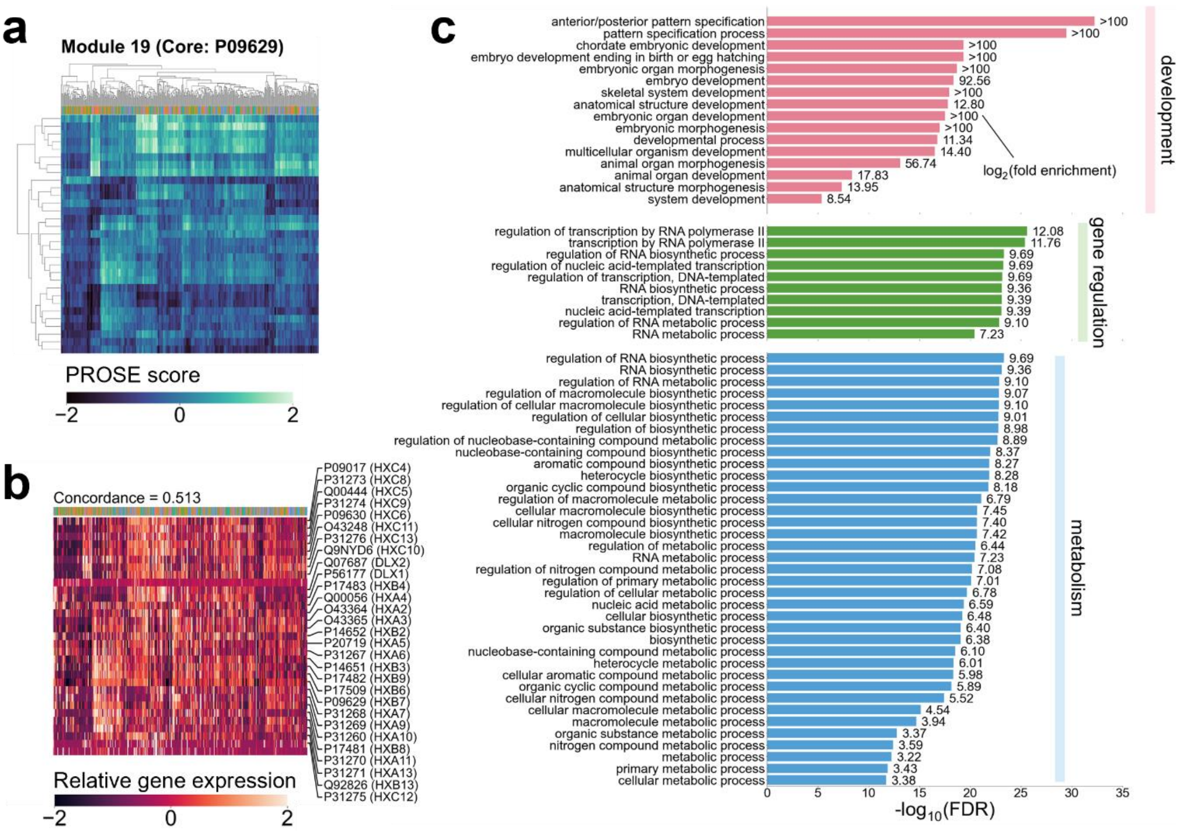


**Supplementary Figure 6. Module 19 containing the HOXA-HOXB locus. a:** PROSE score matrix. **b:** TPM matrix with preserved orderings shows modest visual similarity to the PROSE score matrix. While HOX locus expression is known to be extensively heterogenous between cell lines[3], biclusters are still visually discernible. **c:** Gene set overrepresentation analysis describing the key role of the module in development, gene regulation, and metabolism, as previously described[4]. Significant GO-slim Biological Process terms with FDR < 0.05 by Fisher's exact test are shown. Putative targets of the HOX transcription factors (or co-regulatory elements) can be obtained by gradually decreasing the FastICA-FDR threshold (for instance, from a strict *p*-value of $10^{-6}$ to $10^{-3}$; individual values specified in **Supplementary File 2-5**).

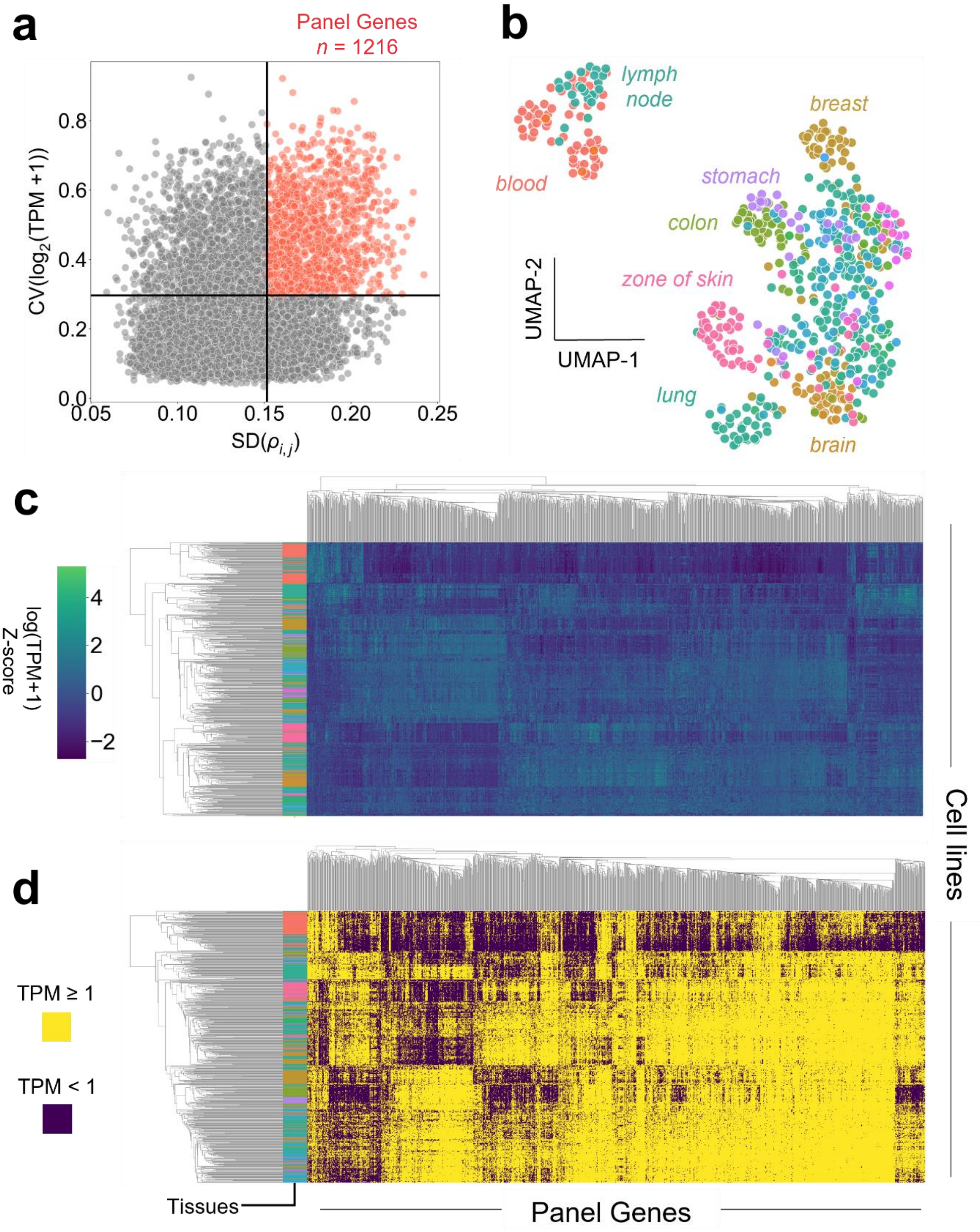


**Supplementary Figure 7. Feature vector (panel gene) selection and characterization. a:** Panel genes identified as the top 25% of genes by both variability in the co-expression matrix (standard deviation of gene-gene Spearman correlations $\rho_{i,j}$) and variability in mRNA expression levels (coefficient of variation; CV; standard deviation divided by the mean of log-transformed TPM). **b:** UMAP constructed from TPM of panel genes demonstrating tissue-specific clustering. Visually prominent clusters were manually annotated. **c:** Heatmaps demonstrating sufficiency of panel genes in capturing differential expression across cell lines **d:** As above, but with binarized mRNA existence.

### Supplementary Table 1. Criteria for protein support level classification

| Protein support type | Criteria |
|---|---|
| Two peptide (unique) support | Supported by at least two uniquely-mapping peptides of FDR < 0.01 |
| One peptide (unique) support | Supported by exactly one uniquely-mapping peptide of FDR < 0.01 |
| Ambiguous support | Supported only by multiply-mapping peptides of FDR < 0.01 |
| No support | Supported only by peptides of FDR > or no peptides detected in the proteomic screen |
| Percolator* | Rescored protein-level FDR < 0.01. N.B. Engine only infers proteins supported by unique peptides. |
| Fido* | Rescored protein-level FDR < 0.01 |
| EPIFANY* | Rescored protein-level FDR < 0.01 |

### Supplementary Table 2. Datasets used in analysis

| | Source/name | Reference | Description |
|---|---|---|---|
| **RNA-seq** | Cancer cell lines | [5] | mRNA-seq data as transcripts per million (TPM) for 622 common cancer cell lines. Used for generating the PROSE feature vector. |
| | CCLE | [6] | mRNA-seq data as TPM for 934 common cancer cell lines. |
| **Proteomics** | Pierce HeLa digest standards | [7] | Pierce HeLa digest standards acquired in quadruplicate via nanoflow LC-MS/MS. DDA mode on an Orbitrap Fusion Lumos Tribrid instrument. Raw data was used. |
| | HeLa lysates | [8] | HeLa lysates acquired via nanoflow LC-MS/MS. DDA mode on a Q Exactive HF instrument. Processed iBAQ data was used. |
| | CCLE | [9][10] | Digests derived from cultured CCLE cell lines, acquired as TMT 10-plexes quantified at the MS3 level. DDA mode on an Orbitrap Fusion or Orbitrap Fusion Lumos instrument. |
| **Other** | DRIVE deep RNAi screen | [11][12] | DEMETER2-corrected dependency scores for the DRIVE deep RNAi screen dataset. |
| **Databases** | COSMIC v94 Sanger cancer gene census (CGC) | [13] | Curated set of known oncogenes, tumor suppressors, fusion and cancer-linked genes. Used for oncogene annotation. |
| | The human transcription factors | [14] | Curated set of known human transcription factors. Used for transcription factor annotation of PROSE-FastICA modules. |
